# Improvement in Outcomes and Prevention of Multiple Organ Dysfunction Syndrome After Cardiac Arrest and Successful Cardiopulmonary Resuscitation by Systemic Administration of Bumetanide in Male Mice

**DOI:** 10.1101/2024.05.08.593259

**Authors:** Zhun Yao, Yuanrui Zhao, Liping Lu, Song Xu, Bin Luo, Jianfei Sun, Ying Zhu, Yanping Wu, Yinping Li, Zhui Yu

## Abstract

**BACKGROUND:** Cardiac arrest (CA) and successful cardiopulmonary resuscitation (CPR) cause post-CA brain injury (PCABI) and extracerebral multiple organ dysfunction (EMOD), leading to low survival and disability in resuscitated patients. The pathogenesis of PCABI is still poorly understood, and no therapeutic-related factors have been identified to improve survival and neurological outcomes to date. Bumetanide is a promising pharmaceutical intervention for some neurological disorders that have some common pathophysiology with PCABI, and it also exhibit systemic protective effects on vital organs under pathological conditions. This study aims to investigate the protective effects of bumetanide on PCABI and EMOD after CA/CPR, and uncover the pathogenesis and biomarkers of PCABI at protein level.

**METHODS:** We generated a hyperkalemia-induced asystole CA/CPR mouse model with bumetanide/vehicle treatment after resuscitation. Survival, neurological outcome, functional outcome, key pathophysiological process underlying PCABI, and injury level of EMOD were evaluated. Proteomics analysis of cerebral cortex was performed for investigating mechanisms of PCABI.

**RESULTS:** Bumetanide significantly improved outcomes after CA/CPR and reduced the main pathophysiological processes of PCABI, including seizures, neurodegeneration, neuroinflammation, decreased cerebral blood flow, blood-brain barrier disruption, and oxidative stress. CA/CPR-induced injury in heart, lung, liver, kidney, spleen, adrenal gland, spinal cord, pennis, and urinary bladder were also alleviated by bumetanide. Proteomic study and experimental verification identified LCN2/NGAL is a potential biomarker for early neuroprognostication and has association with PCABI severity.

**CONCLUSIONS:** Systemic administration of bumetanide improved outcomes and prevented multiple organ dysfunction after CA/CPR. LCN2/NGAL is a novel biomarker for early neuroprognostication at 24 hours after CA/CPR.

**Clinical Perspective:**

1. **What Is New?**
  - Post-resuscitation bumetanide treatment could improve outcomes and prevent brain injury and extracerebral multiple organ dysfunction after hyperkalemia-induced asystole cardiac arrest in mice.
  - Neuronal subpopulations in cerebral cortex were selectively vulnerable, and all major organs had obvious tissue damage with inflammatory cell infiltration and tissue edema after resuscitation.
  - This the first study to identify the pathogenesis and biomarkers of post-cardiac arrest brain injury with proteomic studies, and LCN2/NGAL had a strong and graded association with pathogenesis of PCABI, which could be a marker for monitoring and neuroprognostication.
2. **What Are the Clinical Implications?**
  - Bumetanide may be a novel pharmacological therapy to improve survival and neurological outcomes in post-resuscitation care, and cerebral-specific quantification of LCN2/NGAL could be used for early neuroprognostication at 24 hours after cardiac arrest.
  - Extracerebral multiple organ injury is common and severe after resuscitation, which needs to be given more attention in clinical management.

## Introduction

Cardiac arrest (CA) is the abrupt loss of heart pumping function, resulting in the cessation of systemic blood flow. Return of spontaneous circulation (ROSC) is achieved after successful cardiopulmonary resuscitation (CPR). Even patients successfully get resuscitated form CA, the morbidity and mortality remain significantly high due to a complex combination of pathological processes termed as post-CA syndrome (PCAS).^1^ The four key components of this syndrome comprise post-CA brain injury (PCABI), post-CA myocardial dysfunction (PCAMD), systemic ischemia/reperfusion response (SIRR), and persistent precipitating pathology (PPP) that caused CA and remain unresolved after resuscitation.^1^ PCABI is the leading cause of death after resuscitation, and cardiovascular failure is the main cause of early death.^1–5^ PCABI is a hypoxic ischemic brain injury (HIBI), and pathophysiology of PCABI could be summarized into a “two-hit” model, determined by primary ischemia injury during CA and long-lasting secondary injury following initiation of CPR.^5,6^ Multiple pathophysiological mechanisms contribute to the progression of PCABI, including dysfunction of cerebral oxygen cascade, inflammation, seizures, oxidative stress, cerebral edema, brain tissue edema, blood-brain barrier (BBB) disruption, microvascular dysfunction, and mitochondrial dysfunction.^6,7^ Exposed to complete whole-body ischemia and reperfusion, post-resuscitation extracerebral multiple organ dysfunction (EMOD) is common and heterogenous, and is significantly associated with mortality and poor neuroprognostication.^8^ Optimize vital organ perfusion, reduce the risk of EMOD, and support organ function are fundamental and effective managements to increase the likelihood of intact neurological survival after resuscitation.^3,9^ To date, no therapeutic-related factors, including targeted temperature management, have shown a clear association with positive outcome after ROSC.^2,3,10^ Additionally, no pharmaceutical interventions have been proved to be effective for improving neurological outcome after CA,^7^ and the molecular mechanisms underlying the pathogenesis of PCABI are still poorly understood, which reminds us further investigating mechanisms of PCABI/PCAS to search new drug targets, as well as identifying novel biomarkers that can monitor the development and predict the outcome of PCABI.

Sodium-potassium-chloride cotransporter 1 (NKCC1) is a member of cation-chloride cotransporters that catalyzes electroneutral symport of 1Na^+^, 1K^+^, 2Cl^−^ across cell membranes in mammals.^11^ NKCC1 has an important role in maintaining intracellular ion homeostasis and consequent cell volume homeostasis, neuronal gamma-aminobutyric acid signaling, secretary epithelial transport, immune function, and cell migration.^11,12^ Altered expression and/or function of NKCC1 has been revealed in some central nervous system (CNS) disorders, including cerebral edema, traumatic brain injury, ischemic stroke, neonatal seizures, hydrocephalus, spinal cord injury, neuropathic pain, and other neurological disorders,^12–14^ which have some common pathophysiological mechanisms with PCABI. Thus, NKCC1 is considered as an attractive CNS drug targets in numerous neurological disorders,^12,14–16^ and targeting NKCC1 may also help to improve neurological outcomes after CA. Function of NKCC1 could be specifically inhibited by antagonist bumetanide, which is an FDA-approved loop diuretic that also inhibits NKCC2, an isoform of NKCC1 predominantly expressed in kidney, to modulate water and salt reabsorption.^15,17^ Bumetanide has been used in several clinical studies on a theoretical basis that dysregulation and/or increased expression of NKCC1 contributes to the progression of some neurological disorders.^13,14,17^ Bumetanide is also considered to have systemic protective effects under some pathophysiological conditions, including managing edematous conditions and fluid overload, preventing acute renal insufficiency, treating acute cardiac dysfunction, attenuating acute lung injury, and reducing inflammation.^12,18–21^ Therefore, treatment with bumetanide may also be beneficial to prevent and/or alleviate the multiple organ dysfunction syndrome (MODS) in PCAS.

Here, we report using a hyperkalemia-induced asystole CA/CPR mouse model to duplicate post-resuscitation MODS, including PCABI and EMOD. Histological injury with systemic inflammation and tissue edema, even acute organ failure, could be found in major organ systems after CA/CPR. Systemic administration of bumetanide after resuscitation dramatically improved outcomes and prevented MODS. Expression of NKCC1, the CNS target of bumetanide, was significantly increased in cerebral cortex at the immediate phase of PCAS. We also investigated the molecular mechanisms of PCABI progression on a protein level with proteomics analysis, from which the pathogenesis and biomarkers could be identified. Consistent biomarkers with human serum proteomics studies at the same time point were confirmed and validated by our pre-clinical studies.

## Methods

### Animals

All animal procedures were approved by the Institutional Animal Care and Use Committee of Renmin Hospital of Wuhan University (No. WDRM 20171204). All experiments followed the Animal Research: Reporting In Vivo Experiments (ARRIVE) guidelines 2.0.^22^ Only male mice were used. Wild-type C57BL/6 outbred mice (8 weeks old) were purchased from China Three Gorges University. The mice were housed in a specific pathogen-free room in the Animal Experiment Center of Renmin Hospital of Wuhan University, with a temperature/humidity-controlled environment on a 12-h light/12-h dark cycle and free access of food and water.

### Mouse model of CA and CPR

Mouse CA/CPR model was performed as previously described with minor modifications (Figure 1A).^23^ Briefly, mice were anesthetized by 5% isoflurane, endotracheally intubated, and mechanically ventilated (VentStar, RWD). Mice were maintained on 1.5% isoflurane before CA induction. A thermostat, containing rectal temperature probe and heating pad (RWD), was used to monitor core body temperature and maintained at 37±0.2°C before and during CA. Electrocardiogram was continuously recorded with three subcutaneous needle electrodes (LabChart). A tail-cuff system was placed on the tail for noninvasive blood pressure measurement. A central venous catheterization (CVC, BD) was inserted into the right subclavian vein for blood drainage/transfusion and drug delivery, which enables reach of central circulation faster. After physiological stabilization, 100 µL normal saline containing 5 units of heparin was infused and 200 µL blood was drained through CVC. Asystole CA was induced by rapid infusion of KCl (30 µL, 0.5 M) and confirmed by ECG manifestations. Mechanical ventilation was terminated immediately after CA, and slow autotransfusion started at 3 min after CA. After 9 min of CA, resuscitation was initiated with mechanical ventilation (100% oxygen) and rapid finger chest compressions. A bolus of epinephrine (100 µL, 32 µg/mL) was given at 30 s before CPR, followed by continuous infusion (≤200 µL, 32 µg/mL, 20 µL/min). Body temperature was allowed to spontaneously decrease during CPR and was maintained at 32.0±0.2°C. ROSC was defined as the return of sinus rhythm with normal mean arterial pressure lasting at least 1 minute. Return of spontaneous breathing was synchronously confirmed by presence of respiratory behavior and respiratory wave on ventilator. Mice were weaned from mechanical ventilation and extubated after a spontaneous breathing test. Surgical wound was sutured and disinfected, antibiotic ointment was used for infection prevention and local anesthetic was used for pain relief during recovery period. Mice were returned to cages with easy access of water and feed after 2 h of intensive care in a warm chamber. Post-resuscitation care with closing monitoring was standardized in each mouse. Sham-operated mice were subjected the same surgical procedures but not CA/CPR. Mice failed to achieve ROSC within 4 min were excluded from study.

**Figure 1.**
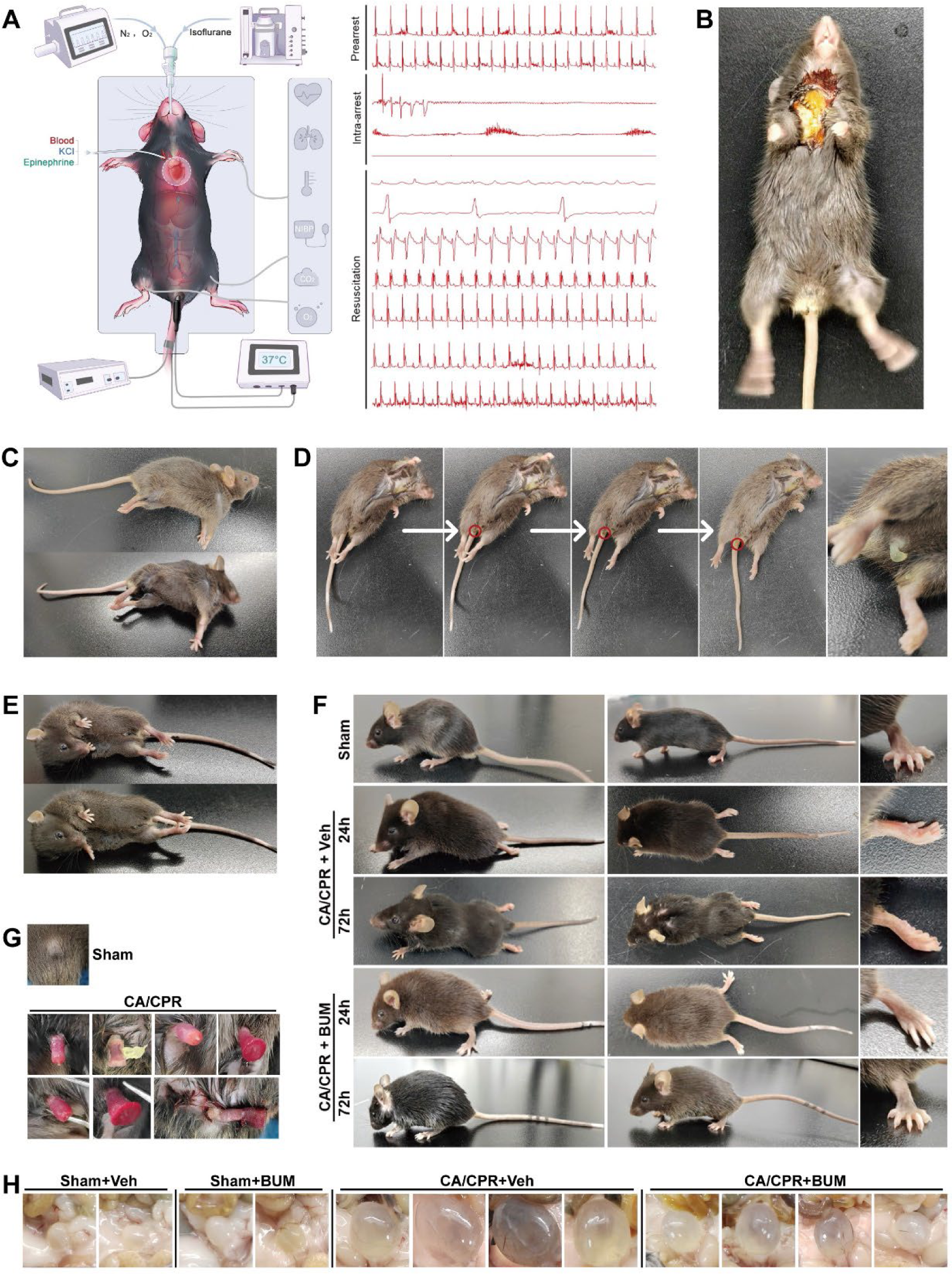
CA/CPR caused severe central nervous system injury. **A**, Schematic diagram of the mouse CA/CPR model and representative electrocardiograph of each stage. **B**, Representative photograph of clinical seizures/myoclonus at 1 hour after ROSC. **C**, Representative photograph of decerebrate rigidity at 6 hours after ROSC. **D**, Representative photographs of paroxysmal hypersympathetic syndrome-like behavior at 3 hours after ROSC in response to external stimuli. Mice showed signs of dysmyotonia and mixed sympathetic hyperactivity (increased heart rate and breathing rate) and parasympathetic (defecation in red circle and ejaculation) hyperactivity. **E**, Generalized tonic–clonic seizures at 12 hours after ROSC. **F**, Hind limb paralysis and body weight loss after CA/CPR. **G**, Priapism and injured penis after CA/CPR. **H**, Urinary retention caused by CA/CPR and alleviated by BUM treatment.

### Randomization, intervention, and blinding

Immediately after resuscitation, mice underwent randomization, generated using a random-number generator, to one of two interventions in a 1:1 ratio: vehicle (Veh, 2% DMSO + 40% PEG300 + 5% Tween-80 + 53% normal saline, 200 µL) or bumetanide (BUM, 2 mg/kg, HY-17468, MedChemExpress), both are given by intraperitoneal injection (i.p.). Bumetanide was dissolved in the vehicle solution with a concentration of 1 mg/ml. The 2 mg/kg i.p. dose of bumetanide was a relatively small dose which could reach minimal NKCC1 inhibitory concentration in CNS.^15^ The urination and body weight loss were closely monitored, subcutaneous injection of 500 µL warm saline was performed when finding a loss of 1 g weight with increased urine output.

The investigators who carried out subsequent experiments were blinded from group assignment. During the analysis phase, the statisticians and analysts were unaware of the group assignment, which were identified as Y and Z.

### Sample size calculation, survival study, and assessment of neurological function and motor function

Based on our previous experience, the three-day survival rate was expected as 20% in CA/CPR mice and 2/3 (67%) in bumetanide-treated CA/CPR mice. Thus, the baseline conversion rate was 20% and absolute minimum detectable effect was 47%, along with the setting values that significance level (α) is 5% and statistical power (1-β) is 80% (two-sided). After calculation, we estimated a sample size of 12 in each group. Survival was recorded every 12 hours for 4 days after CA/CPR. Kaplan-Meier estimates of the survival curve were obtained from each group, and log-rank test was used to perform statistical comparisons. The Cox proportional hazards models were used to study differences in survival.

Neurological function and motor function at 24 hours and 72 hours after CA/CPR were assessed with previously-reported scoring systems^24^ by 2 investigators blinded to operation and treatment. The neurological function scoring system has 6 parameters: level of conscious, corneal reflex, respirations, righting reflex, coordination, and movement/activity. Each parameter was scored as 0 (impaired), 1 (mildly impaired), or 2 (normal). The total score was reported as the neurological function score. Dead mice were scored at 0 points and were excluded from statistical analysis.

A 9-point scoring system was used to evaluate general motor function reflecting overall neurological deficits.^25^ Mice were placed on a rope, horizontal bar, and vertical screen, and its performance was scored as 0 to 3 respectively. The total score was computed (9 points=normal and 0 points = severe injury). A 4-point scoring system was used to evaluate the motor function of hind limbs (0 points = complete paraplegia and 4 points = normal).^26^ Dead mice were excluded from the statistical analysis.

To ensure the accuracy of score, two investigators independently assessed the neurological function score and motor function score, and all discrepancies were resolved with additional assessment by a third investigator. The neurological function score and motor function score were compared with Mann–Whitney U test.

### Electrocardiogram study, core body temperature measurement, body weight loss, and exploratory laparotomy

Mice were anesthetized, intubated, ventilated, and maintained on 1.5% isoflurane before sacrificed. Electrocardiogram study and core body temperature measurement were performed as described above. Body weight was measured, and loss of which was calculated. The abdomen was opened for exploratory laparotomy, and representative images of whole abdominal organs were taken.

### Histology and staining

Detailed descriptions of reagents and the protocols for are provided in the Supplementary Methods.

### Western blotting

A standard protocol was used for Westen blotting.^27^ Detailed protocols are provided in the Supplementary Methods.

### Evaluation of BBB permeability

Evans blue (EB, 2% in normal saline, 4 mL/kg, HY-B1102, MedChemExpress) extravasation assay was used for quantifying BBB permeability. Detailed protocols are provided in the Supplementary Methods.

### Measurement of cerebral blood flow

Laser speckle contrast imaging (LSCI, Moor Instruments) was used for monitoring global cerebral blood flow (CBF), a key component of brain hemodynamics. Detailed protocols are provided in the Supplementary Methods.

### Quantitative data-independent acquisition proteomic analysis

A total of 10 samples of mice brain cortex at 24 h after operation were collected, and 4 samples from age/weight-matched healthy mice were selected as control. We collected 6 samples in the CA/CPR group, which were equally divided into 2 subgroups: CA1 and CA2. CA1 represents the severe PCABI with poor prognosis, and CA2 represents relatively good outcome. The criteria for grouping were based on brainstems reflexes and the responses to pain, which are the most useful indicators of outcome in neuroprognostication and are reflected by corneal reflex and level of conscious (response to tail pinch) in the neurological function score. Thus, we selected mouse had impaired/mild impaired (0 or 1 score) corneal reflex and conscious level, along with the absence of righting reflex (0 score), and grouped into CA1. Similarly, we selected mouse had mild impaired/normal (1 or 2 score), along with the presence of righting reflex (1 or 2 score), and grouped in to CA2.

Data-independent acquisition (DIA) proteomic analysis and subsequent bioinformatic analysis, including GO (gene ontology), KEGG, COG (cluster ortholog groups of proteins) annotation, and PPI (protein-protein interaction) analysis, of the differentially expressed proteins (DEPs) were performed. Detailed descriptions are provided in the Supplement Methods.

### Assessment of mesenteric prefusion and intestinal microcirculation

To directly visualize the mesenteric perfusion and intestinal microvascular perfusion, a non-invasive hand-held sidestream dark field (SDF) microscope (V100, Medsoft System, Guangzhou, China) was used. The most vulnerable intestinal section was externalized through exploratory laparotomy, and images, covering an area approximately 2.5 cm^2^, of corresponding supplying mesenteric vessels and intestinal microvascular tissues were acquired according to the manufacturer’s instructions by a well-trained researcher. The results were also analyzed with the same system.

### Statistical Analysis

Data of continuous variables were presented as mean ± standard error of the mean (SEM) or medians with interquartile range. Unpaired two-tailed Student’s t test was used for comparing data of two groups. One-way analysis of variance (ANOVA) followed by Sidak’s corrections with post hoc comparisons was used for multigroup comparisons. A value of P<0.05 was considered statistically significant. All analyses were performed with the use of GraphPad Prism, version 8.0 (GraphPad) and SPSS, version 27.0 (IBM SPSS).

## Results

### CA/CPR caused severe CNS injury and systemic administration of bumetanide improved survival, neurological outcome, and functional outcome after CA/CPR

There was no significant difference in the perioperative baseline characteristics between CA/CPR mice treated with vehicle or bumetanide (Table S1). On achieving ROSC, all mice were unresponsive, which is a marker of significant brain injury. In the vehicle-treated CA/CPR group, mice showed significant signs of HIBI, including clinical seizures/myoclonus (Figure 1B, Video S1 and S2), decerebrate rigidity (Figure 1C), paroxysmal hypersympathetic syndrome-like behavior (or mixed parasympathetic and sympathetic hyperactivity, often induced by external stimuli, Figure 1D), as well as generalized tonic–clonic seizures (Figure 1E) which was significantly associated with poor outcome and might lead to recurrent arrest as we observed. CA/CPR also induced other signs of brain/spinal injury, including hind limb paralysis (Figure 1F), rotational behavior (Video S3), priapism (Figure 1G and S1A), and urinary retention (Figure 1H). Given that there were no signs of urinary obstruction (all mice experienced urinary incontinence after induction of CA during operation and had micturition in the first 3 hours after resuscitation, Figure S1B), we thought that urinary retention was caused by the nerve injury in brain/spinal cord that disrupted messaging between the brain and bladder. On account of the absence of sexual stimulation, the priapism maybe a sign of spinal cord injury. Notably, we observed that the priapism was usually painful and extremely hard, mice may even injured penis themselves, which suggests an ischemic (low-flow) subtype (Figure 1G and S1A). We also found the faeces were black, dry, and could not be fully discharged after CA/CPR (Figure S1A). We also observed mice licked their ventral hair more frequently and make it soaked after CA/CPR, which may also be a manifestation of CNS injury, while treatment with bumetanide decreased this abnormal behavior (Figure S1A).

Treatment with bumetanide after ROSC dramatically improved survival (Figure 2A). The 12-point neurological function score (Figure 2B and 2C), 9-point motor score (Figure 2D and 2E), and 4-point hind-limbs motor score (Figure 2F and 2G) were also significantly higher in bumetanide-treated mice than in vehicle-treated mice at 24 hours and 96 hours after CA/CPR. Administration of bumetanide also decreased the body weight loss after CA/CPR (Figure 2H). The core body temperature was also higher in the bumetanide-treated mice than the vehicle-treated mice at 24 hours after ROSC (Figure 2I).

**Figure 2.**
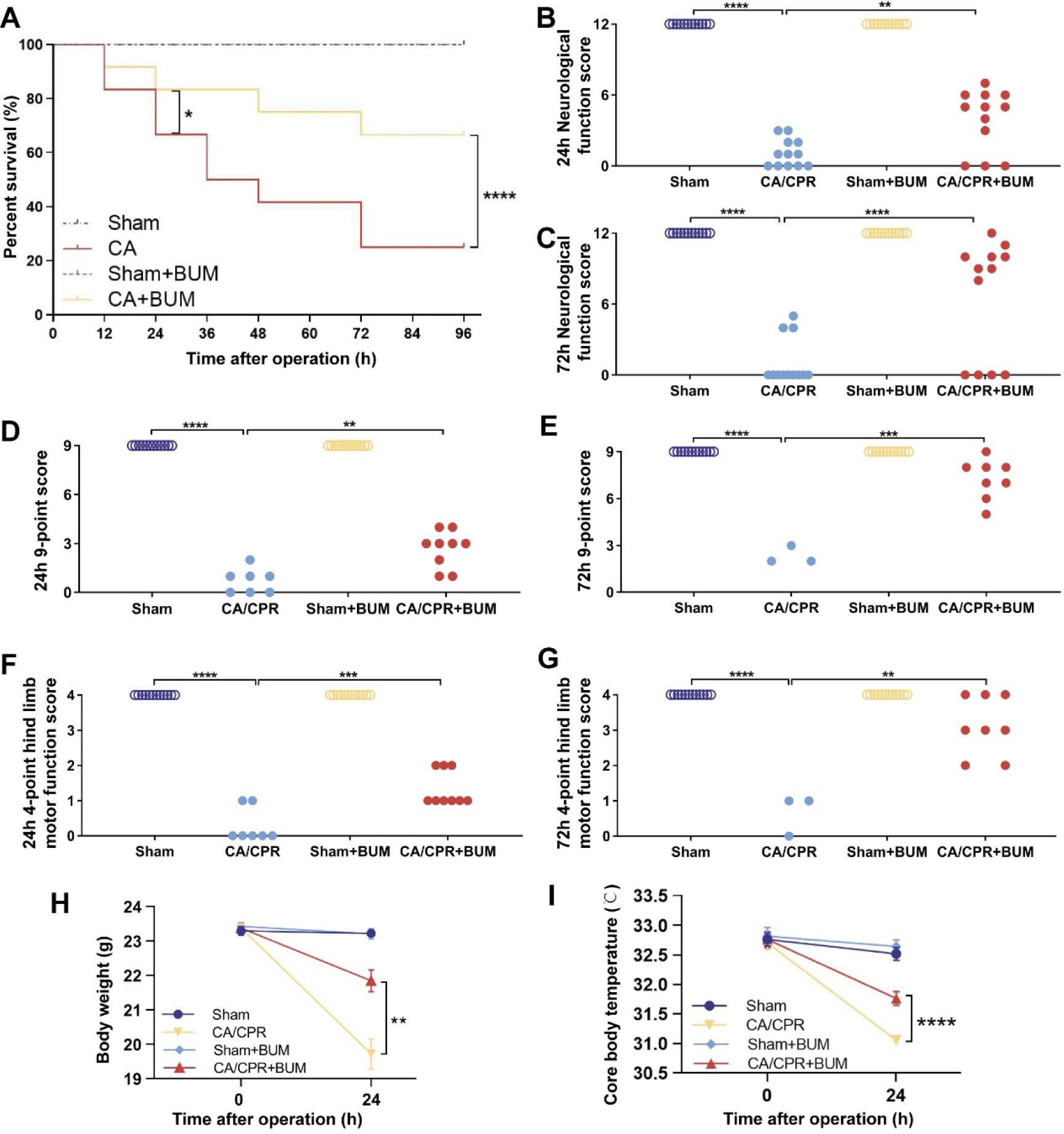
Bumetanide improved survival, neurological outcome, and functional outcome after CA/CPR. **A**, Kaplan–Meier analyses of cumulative survival during the first 4 days after CA/CPR. Neurological function score at 24 hours (**B**) and 72 hours (**C**) after operation. Dead mice were scored 0. (n=12) 9-point function score at 24 hours (**D**) and 72 hours (**E**) after operation. 4-point hind limb motor function score at 24 hours (**F**) and 72 hours (**G**) after operation. Body weight loss (**H**) and core body temperature (**I**) at 24 hours after operation. Data are presented as mean ± SEM. Dead mice were excluded from the statistical analysis. \**P* < 0.05, \*\**P* < 0.01, \*\*\**P* < 0.001, **** *P* < 0.0001.

### BUM prevented histological injury and reduced neuroinflammation in brain and spinal cord after CA/CPR

To evaluate histopathologic changes in brain and spinal cord, we performed HE staining and Fluoro-Jade B (FJB) staining. In FJB-staining brain slides, CA/CPR significantly increased the ratio of FJB-positive cells (Figure 3A). Additionally, we identified neuronal subpopulations in cerebral cortex (especially piriform cortex), amygdaloid nucleus, hypothalamus, corpus striatum, but not hippocampus, thalamus, cerebellum, or medulla oblongata, were selectively vulnerable (Figure 3B). In HE-staining brain slides, CA/CPR induced selective eosinophilic neuronal death (SEND), characterized by deeply stained red, pycnotic neurons and shrunken nucleus, vacuolization, and tissue edema (Figure 3C). In the spinal cord slides, CA/CPR also induced extensive SEND, vacuolization, tissue edema, and neurodegeneration (Figure 3C). Treatment with bumetanide significantly deceased numbers of SEND neurons, numbers of vacuoles, severity of edema, and numbers of FJB-positive cells in brain and spinal cord (Figure 3A through 3C).

**Figure 3.**
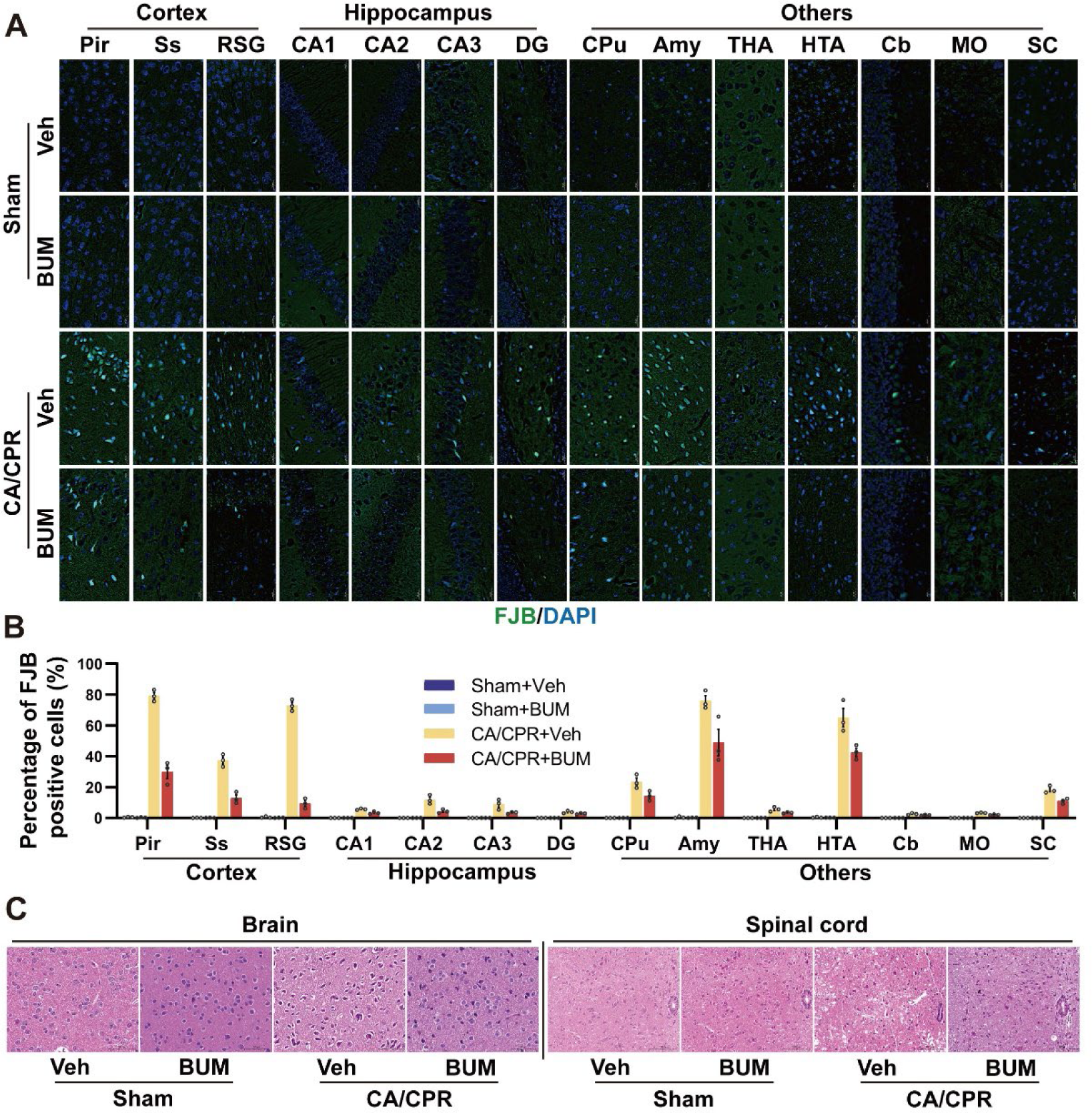
Bumetanide alleviated neurodegeneration and histological injury in central nervous system after CA/CPR. **A**, Representative photomicrographs of FJB (green)/DAPI (blue)-staining brain sections and spinal cord sections at 72 hours after CA/CPR. Neuronal subpopulations in piriform cortex (Pir), somatosensory cortex (Ss), retrosplenial granular cortex (RSG), CA1-3, dentate gyrus (DG), caudate putamen (CPu), amygdaloid nucleus (Amy), thalamus (THA), hypothalamus (HTA), cerebellum (Cb), medulla oblongata (MO), and spinal cord (SC) were examined. **B**, Percentage of FJB-positive cells in brain and spinal cord. (n=4) Data are presented as mean±SEM. \**P* < 0.05, \*\**P* < 0.01. **C**, Representative photomicrographs of HE-staining brain sections and spinal cord sections at 72 hours after CA/CPR. Bumetanide alleviated CA/CPR-caused selective eosinophilic neuronal death, vacuolization, and tissue edema.

We analyzed the level of neuroinflammation with glial fibrillary acidic protein/ ionized calcium-binding adaptor molecule 1 double-staining. Astrocytes and microglia were markedly activated in CA/CPR mice compared with the sham-operated mice, and treatment with bumetanide after CA/CPR decreased activation of these two glias compared with the vehicle-treated group (Figure 4A through 4C).

**Figure 4.**
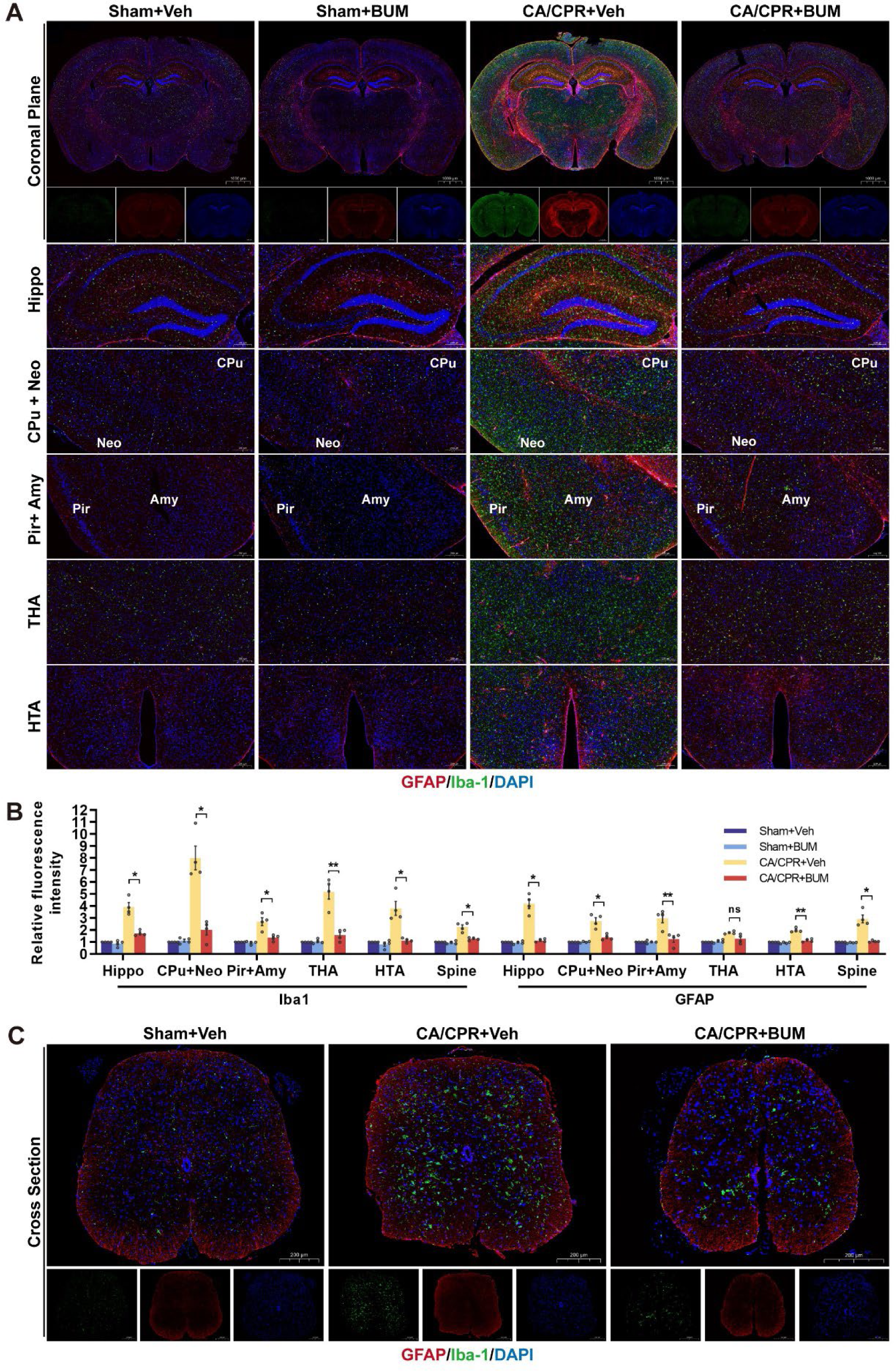
Bumetanide reduced neuroinflammation in brain and spinal cord after CA/CPR. **A**, Representative photomicrographs of GFAP (red) /Iba-1 (green)/DAPI (blue)-staining brain sections at 24 hours after CA/CPR. Brain regions of hippocampus (Hippo), caudate putamen (CPu), neocortex (Neo), piriform cortex (Pir), amygdaloid nucleus (Amy), thalamus (THA), and hypothalamus (HTA) were examined. CA/CPR induced severe neuroinflammation in whole brain, and bumetanide treatment alleviated neuroinflammation. **B**, Relative immunofluorescence signal intensity of GFAP/Iba-1 in different brain regions and spinal cord. (n=4) Data are presented as mean±SEM. \**P* < 0.05, \*\**P* < 0.01; ns, not significant. **C**, Representative photomicrographs of GFAP (red) /Iba-1 (green)/DAPI (blue)-staining spinal cord sections at 24 hours after CA/CPR.

### BUM improved CBF, prevented BBB disruption, and reduced oxidative stress after CA/CPR

To evaluate the alternations of CBF, which is a key component of brain hemodynamics and affects the convective oxygen/glucose delivery for cerebral metabolism, LSCI was performed at 24 hours after ROSC. There was no significant difference of CBF between vehicle/bumetanide-treated sham-operated mice (Figure 5A). CA/CPR caused marked decrease of CBF compared with sham, and treatment with bumetanide improved CBF after resuscitation (Figure 5A and 5B).

**Figure 5.**
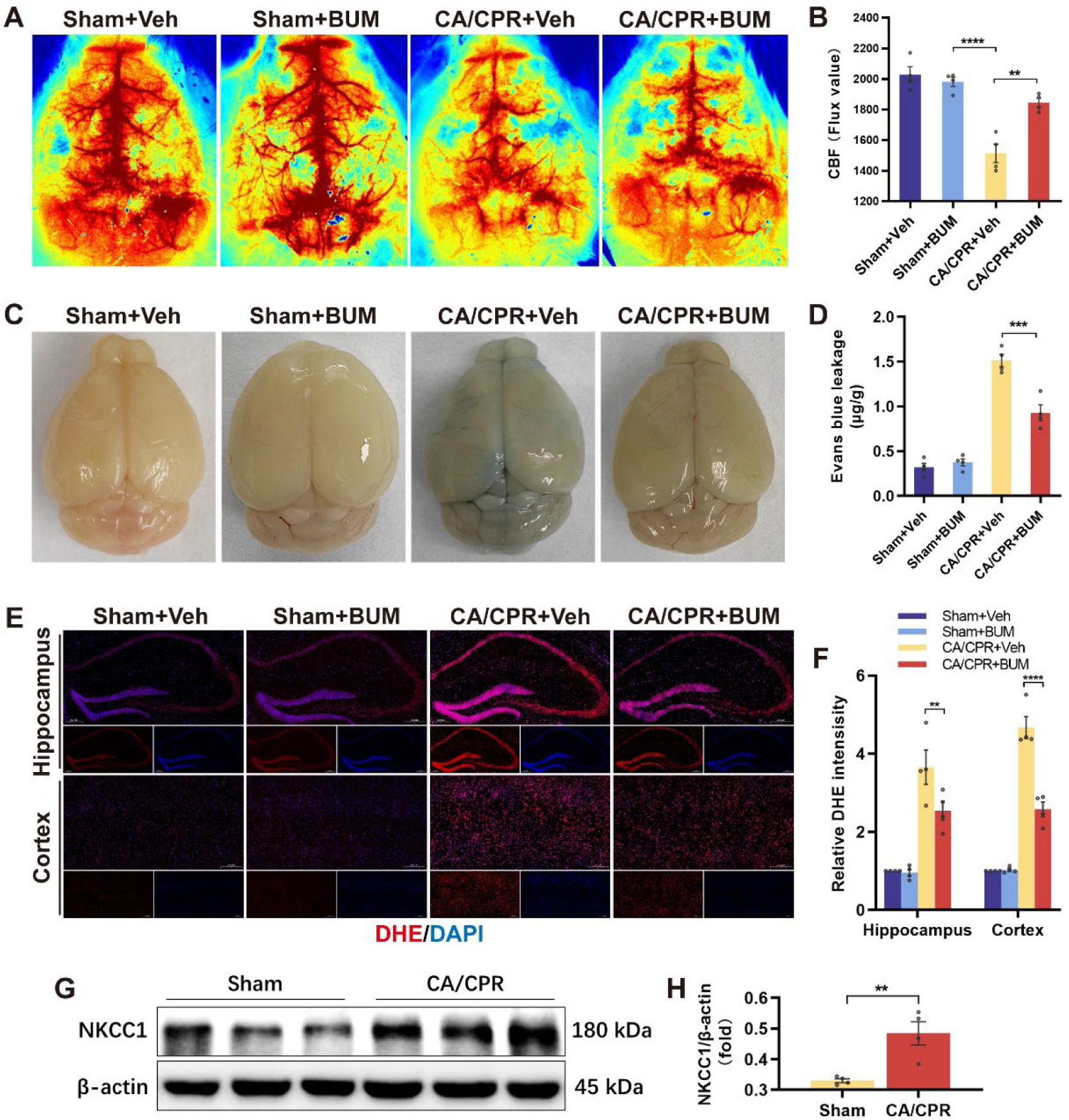
Bumetanide prevented multiple pathophysiological process of brain injury after CA/CPR. **A**, Representative laser speckle contrast images showing reduced cortical cerebral blood flow (CBF) at 24 hours after CA/CPR model, and treatment with bumetanide prevented the reduction of CBF. **B**, Quantitative graph of CBF using the Flux value. (n=3) **C**, Representative whole brain photographs after Evans blue injection at 24 hours after operation. **D**, Quantitative graph of Evan blue leakage in brain. (n=3) **E**, Representative photomicrographs of dihydroethidium (DHE)-staining brain sections in hippocampus and cortex at 24 hours after operation. **F**, Relative immunofluorescence signal intensity of DHE in hippocampus and cortex. (n=3) **G**, Western blot analysis of NKCC1 in the cerebral cortex at 1 hour after sham-operation or CA/CPR. **H**, Quantitative graph of the ratio between NKCC1 and β-actin. Data are presented as mean±SEM. \*\**P* < 0.01, \*\*\**P* < 0.001, \*\*\*\**P* < 0.0001.

BBB disruption and oxidative stress are key components of mechanisms upon PCABI. Intravenous injection of EB was used to detect BBB integrity, and CA/CPR increased the BBB permeability with more EB extravasation compared with sham, while treatment with bumetanide after CA/CPR alleviated BBB disruption as evidenced by less EB extravasation compared with treatment with vehicle (Figure 5C and 5D). To evaluate the oxidative stress level, we detected the reactive oxide species (ROS) production with dihydroethidium staining. CA/CPR increased the production of ROS, which was mitigated by bumetanide treatment (Figure 5E and 5F). These results suggested that bumetanide played a neuroprotective role after CA/CPR by improving CBF, preventing BBB disruption, and reducing oxidative stress.

### The expression of NKCC1 in cerebral cortex significantly increased at the immediate phase after CA/CPR

To examine the expression level of NKCC1, the CNS target of bumetanide, we performed western blotting to quantify the expression change. We found that a significantly increased expression of NKCC1 in cerebral cortex at 1 hour after ROSC (Figure 5G and 5H), which indicated a novel molecular mechanism mediated the progression of PCABI at the immediate phase in PCAS.

### Cerebral cortex proteomics identified biomarkers and pathogenesis of PCABI at 24h after ROSC

To directly investigate the pathogenesis and biomarkers of PCABI, we performed cerebral cortex DIA proteomics at 24 hours after ROSC to study the molecular processes underlying injury and identify potential treatment targets and novel biomarkers (Excel file S1). Comparisons between CA and control represents the proteomic alternations after CA/CPR, and comparison between CA2 and CA1 represents the proteomic alternations according to the neurological outcome or the severity of PCABI.

A total of 7,745 proteins and 85,341 protein precursors were identified (Figure 6A). The quality control analysis of proteomics experiments indicated our quantification results were highly reliable and reproducible at protein level (Figure 6B and S2A through S2E). We identified 118 DEPs in CA1/control, 93 DEPs in CA2/control, and 22 DEPs in CA2/CA1 (Figure 6C). These results reflected a stronger proteomic response in the poor neuroprognostication than the relatively good neuroprognostication. The DEPs were analyzed by the hierarchical clustering method and the results visualized in the heatmap (Figure 6D).

**Figure 6.**
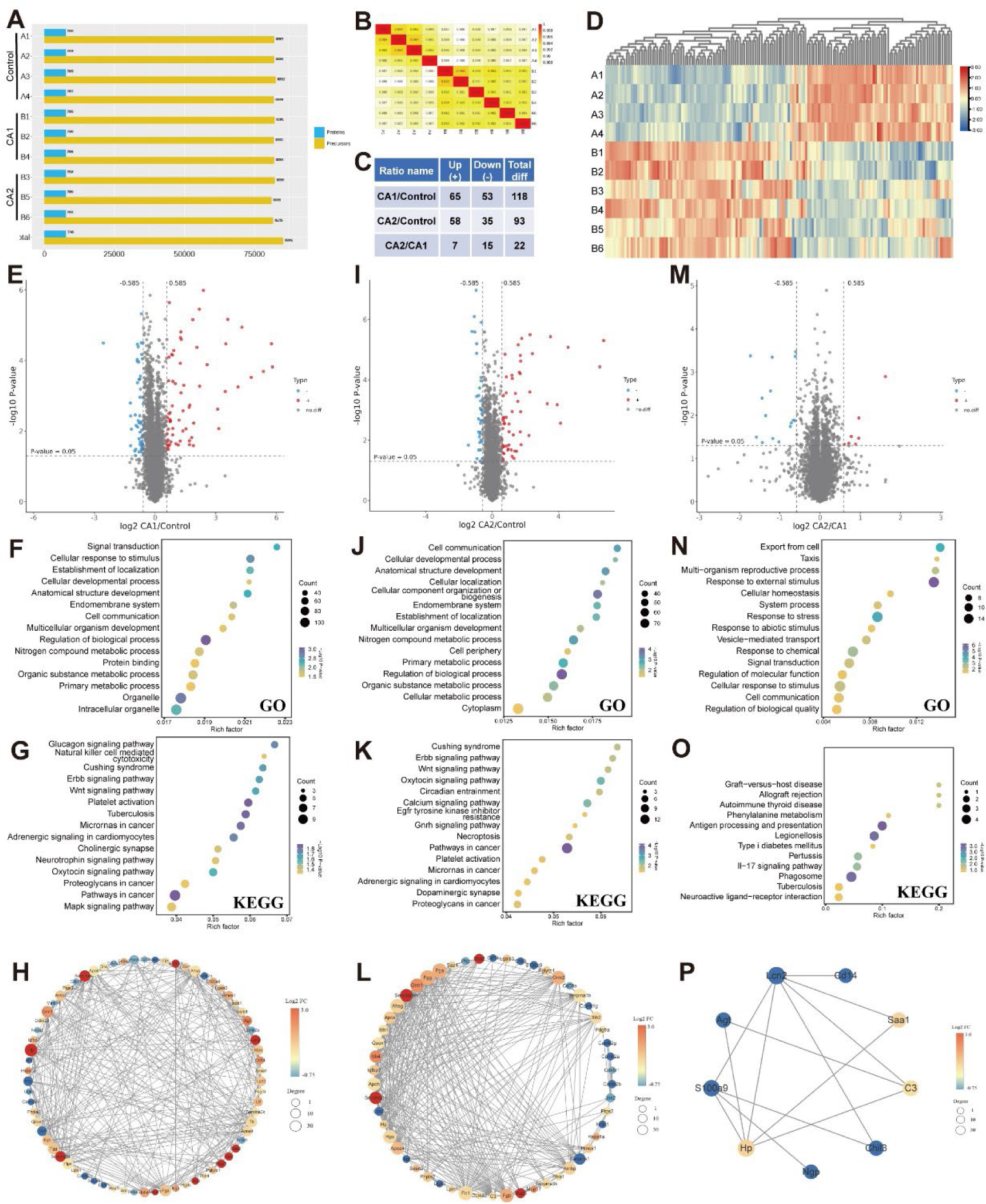
Proteomic profiling of the cerebral cortex and bioinformatics analysis of the differentially expressed proteins (DEPs). Three subgroups were set in this proteomic study: control (n=4, A1-4, healthy mice), CA1 (n=3, B1, B2, B4, poor neuroprognostication mice after CA/CPR), CA2 (n=3, B3, B5, B6, relatively good neuroprognostication mice after CA/CPR). Cerebral cortex was collected at 24 hours after CA/CPR. **A**, The numbers of quantified peptides and proteins in the samples. **B**, Pearson’s correlation of protein quantitation. **C**, Summary of the DEPs in the comparison of three subgroups. **D**, Heat map of all DEPs. The volcano (**E**), GO-based enrichment analysis (**F**), KEGG-based enrichment analysis (**G**), and PPI network (**H**) of DEPs in the comparison of CA1/control are shown in the left. The volcano (**I**), GO-based enrichment analysis (**J**), KEGG-based enrichment analysis (**K**), and PPI network (**L**) of DEPs in the comparison of CA2/control are shown in the middle. The volcano (**M**), GO-based enrichment analysis (**N**), KEGG-based enrichment analysis (**O**), and PPI network (**P**) of DEPs in the comparison of CA2/CA1 are shown in the right.

The volcano plots, GO, and KEGG annotation and enrichment analysis, and PPI networks of DEPs in CA1/control (Figure 6E through 6H), CA2/control (Figure 6I through 6L), and CA2/CA1 (Figure 6M through 6P) are shown. GO and KEGG enrichment analysis indicated that DEPs mainly participated in the following activities and pathways: signal transduction, cellular response to stimulus, regulation of biological process, organelle, natural killer cell mediated cytotoxicity, Cushing syndrome, nitrogen compound metabolic process, Erbb, Wnt, oxytocin, glucagon and MAPK signaling pathway, platelet activation in CA1/control; cell communication, cellular component organization or biogenesis, endomembrane system, nitrogen compound metabolic process, regulation of biological process, cellular metabolic process, cytoplasm, Cushing syndrome, Erbb, Wnt, calcium, oxytocin, and GnRH signaling pathway, and platelet activation in CA2/control; response to external stimulus, stress, and chemical, signal transduction, cellular response to stimulus, cell communication, antigen processing and presentation, IL-17 signaling pathway, phagosome, and Legionellosis in CA2/CA1 (Figure 6F, 6G, 6J, 6K, 6N, 6O, and S3A through 3C). The COG groups of proteins annotation were performed, and signal transduction mechanisms, posttranslational modification, protein turnover, chaperones, and defense mechanisms were identified (Figure S4A through S4C). These results indicated that an infection complication and stress response occurred, and inflammation and immune response played an important role in the progression of PCABI.

We identified 60 common DEPs between CA1/control and CA2/control, which represented the common proteomic alternations underlying PCABI at 24 hours after ROSC (Figure 7A). Additionally, we identified 10 DEPs from the above results that were also significantly differentiated in CA2/CA1, including serum amyloid A-1, haptoglobin, solute carrier organic anion transporter family member 1A6, chitinase-like protein 3, major urinary protein 3, CD14, angiotensinogen, neutrophil gelatinase-associated lipocalin (NGAL), S100-A9, and heat shock 70 kDa protein 1A, which represented the severity-associated and neuroprognostication-associated candidate biomarkers upon PCABI (Figure 7B). Going a step further, we identified 2 proteins, NGAL and angiotensinogen, that were also significantly differentiated in the comparison between good and poor neuroprognostication in human serum proteomic studies at 24 hours after ROSC.^28^ The expression level and spatial expression characteristics of NGAL (Figure 6C through 6E) and CD14 (Figure S5) in brain were examined by immunofluorescence. We found the cerebral expression of NGAL was extremely low in the sham-operated mice (Figure 7C), and increased expression were found after CA/CPR (Figure 7D and 7E). Bumetanide treatment reduced the cerebral expression of NGAL (Figure 7D and 7E). Additionally, we compared the NGAL-positive and FJB-positive encephalic region, and found there were highly overlapping areas, which indicated that NGAL was highly correlated with neuronal degeneration, and consequently progression of PCABI (Figure 7D). We also found that the cerebral expression level of CD14 was increased after CA/CPR, and was decreased after bumetanide treatment (Figure S5).

**Figure 7.**
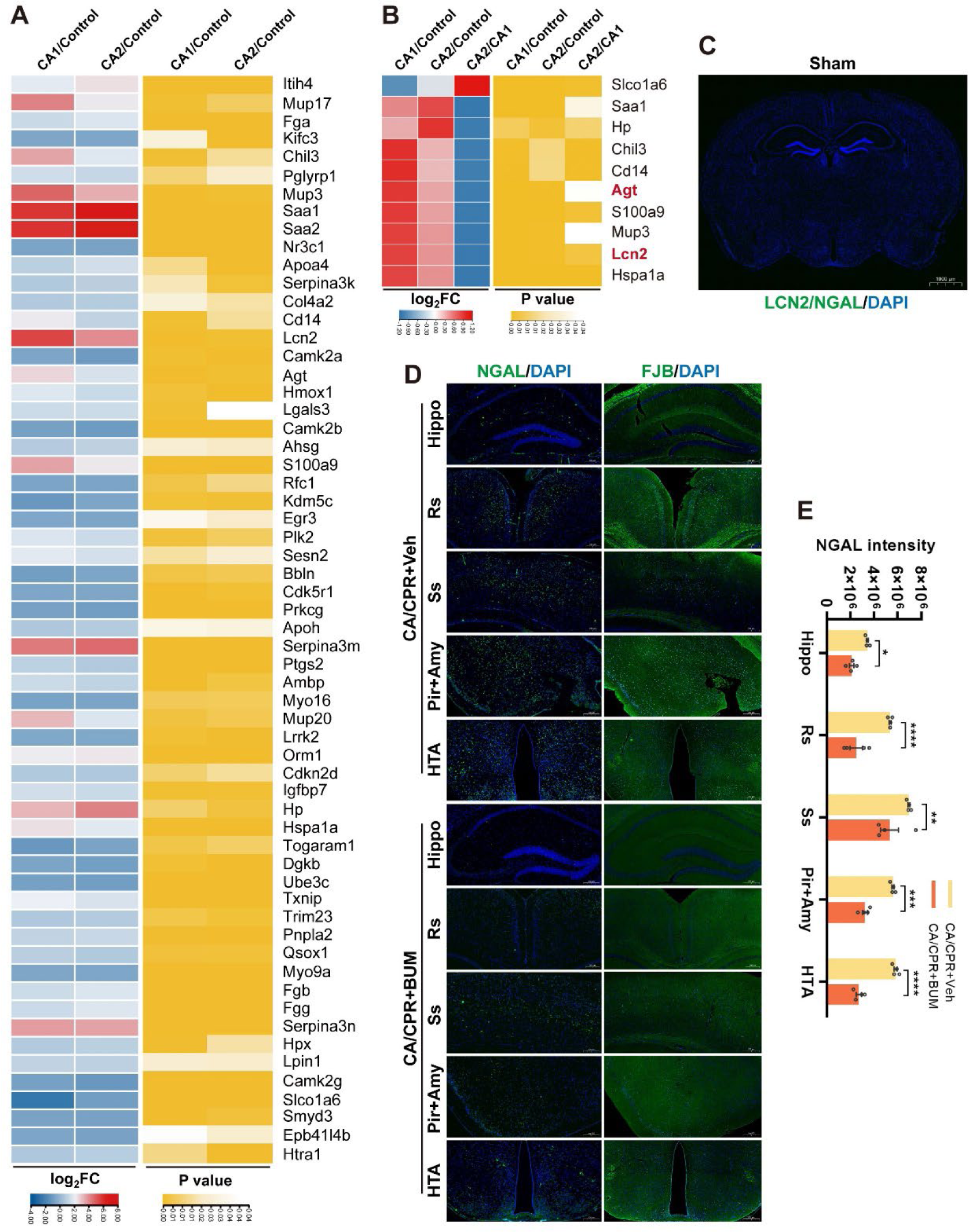
Identification of the pathogenesis and biomarkers of brain injury at 24 hours after CA/CPR. **A**, Common differentially expressed proteins (DEPs) and associated *P* values in CA1/control and CA2/control. **B**, Common DEPs and associated P values in CA1/control, CA2/control, and CA2/CA1. Agt and Lcn2 are also DEPs in the human serum proteomic studies of the neuroprognostication-associated biomarkers at 24 hours after CA/CPR. **C**, Representative photomicrographs of LCN2/NGAL (greeen)/DAPI (blue)-staining brain sections after sham-operation. The expression level of NGAL is extremely low in brain after sham operation. **D**, Representative photomicrographs of LCN2/NGAL (green)/DAPI (blue)-staining and FJB (green)/DAPI (blue)-staining brain sections at 24 hours after CA/CPR with vehicle/bumetanide treatment. Brain regions in hippocampus (Hippo), retrosplenial cortex (Rs), somatosensory cortex (Ss), piriform cortex (Pir), amygdaloid nucleus (Amy), and hypothalamus (HTA) were examined. The NGAL intensity was increased after CA/CPR, and was reduced after bumetanide treatment in brain. There was a considerable overlap between NGAL-positive and FJB-positive areas. **E**, Relative immunofluorescence signal intensity of NGAL in different brain regions. (n=3 or 4) Data are presented as mean±SEM. \**P* < 0.05, \*\**P* < 0.01, \*\*\**P* < 0.001, **** *P* < 0.0001.

### Bumetanide improved electrocardiogram manifestation and ameliorated histological myocardial injury after CA/CPR

To investigate the changes of cardiac functions and myocardial damage, we studied the electrocardiogram (Figure 8A) and performed pathological examination with HE staining and Masson staining (Figure 8B). CA/CPR induced significant changes in electrocardiogram compared with sham, and manifestations of which are abnormal and heterogeneous, including decreased heart rate (Figure 8A and 8C). Histopathological examination showed severe myocardial damage induced by CA/CPR at 72 hours after ROSC, including myocytolysis, inflammatory cell infiltration, fragmentation of the myocardium, disordered arrangement of myocardium, and increased myocardial fibrosis with increased collagen volume fraction (Figure 8B and 8D). Treatment with bumetanide after resuscitation improved electrocardiogram manifestation, alleviated myocardial damage, and reduced myocardial fibrosis (Figure 8A through 8D).

**Figure 8.**
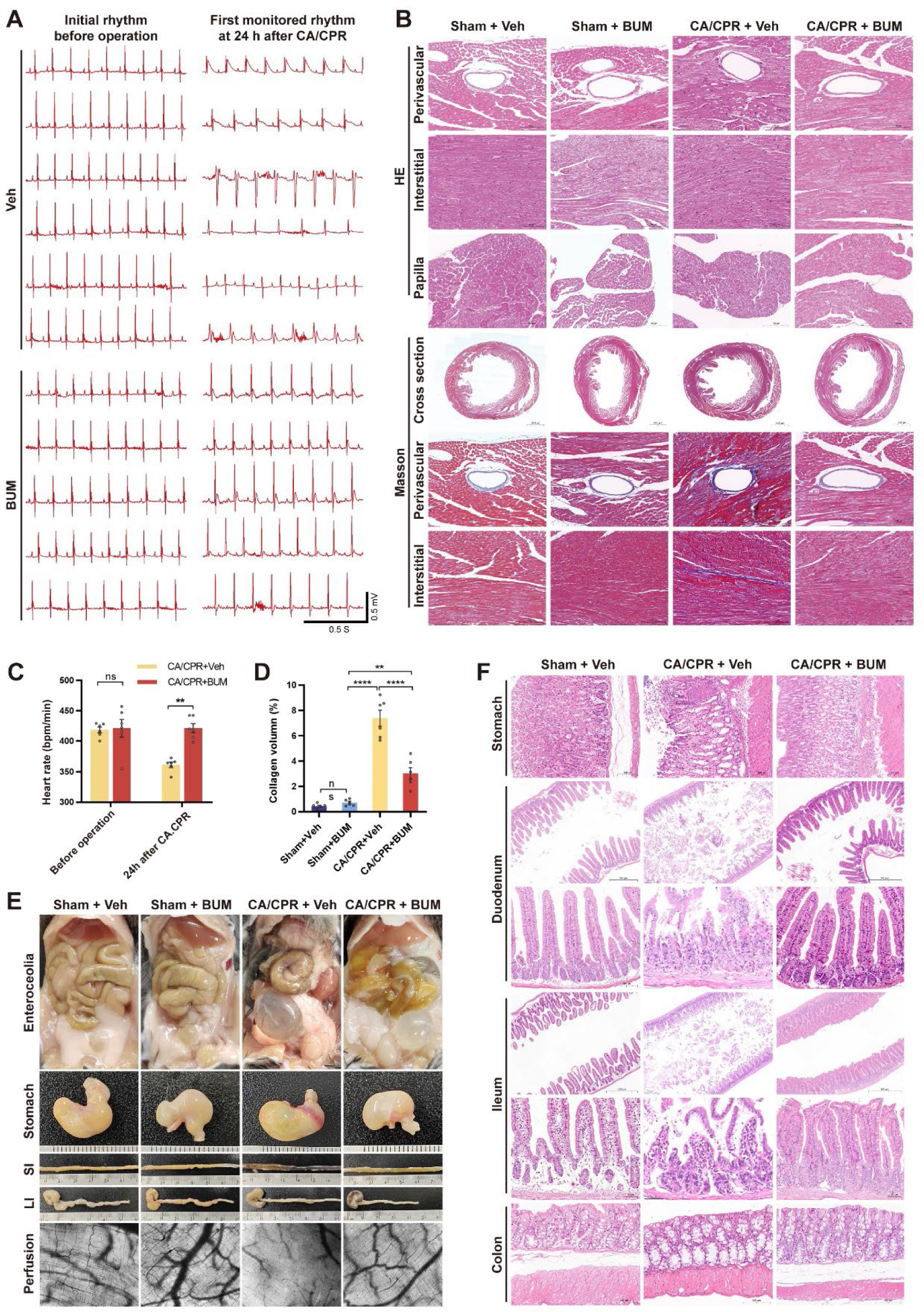
Bumetanide attenuated acute myocardial dysfunction and gastrointestinal injury after CA/CPR. **A**, Representative photographs of electrocardiogram showing initial rhythm before operation and first monitored rhythm at 24 hours after CA/CPR. The electrocardiogram manifestations were abnormal and heterogeneous in the vehicle-treatment CA/CPR mice, and bumetanide improved the electrocardiogram manifestations. **B**, Representative photomicrographs of HE-staining and Masson-staining heart sections 72 hours after CA/CPR. Bumetanide alleviated myocytolysis, inflammatory cell infiltration, fragmentation of the myocardium, disordered arrangement of myocardium, and increased myocardial fibrosis after CA/CPR. Quantitative graph of heart rate (**C**) and collagen volume (**D**) at 24 hours after CA/CPR. (n=6) Data are presented as mean±SEM. \*\**P* < 0.01, \*\*\**P* < 0.001, \*\*\*\**P* < 0.0001; ns, not significant. **E**, Representative photographs of enterocoelia, stomach, small intestine (SI), large intestine (LI), and photomicrographs of intestinal perfusion at 24 hours after CA/CPR. Gastrointestinal injury and urinary retention were found by exploratory laparotomy after CA/CPR. In stomach, obvious hyperemia in the glandular stomach and gases distension were found. In SI, obvious necrosis and intestinal obstruction were found. There were no obvious changes in the LI. Sidestream dark field microscope showed the perfusion characteristics of the most vulnerable segment in SI, indicating no signs of obstruction but significant low-perfusion with dysfunction of microcirculation. Treatment with bumetanide alleviated the above injuries. **F**, Representative photomicrographs of HE-staining stomach, duodenum, ileum, and colon sections at 24 hours after CA/CPR. Bumetanide alleviated tissue edema, inflammatory cell infiltration, ulcerations, and necrosis in the glandular stomach. Erosion was obvious in SI, and severe necrosis was found in ileum. Treatment with bumetanide alleviated the injuries in SI and prevented morphological changes in colon.

### CA/CPR caused severe acute gastrointestinal injury and treatment with bumetanide partly alleviated it

Severe acute gastrointestinal injury (AGI) after CA/CPR was found in exploratory laparotomy (Figure 8E). At 1 hour after ROSC, we found hyperemia in the glandular stomach (but not the forestomach), small intestine (duodenum, jejunum, ileum), except for the large intestine (Figure S6A through S6D). We noticed that the most vulnerable intestinal section was the middle of the small intestine with obvious hemorrhage (Figure S6E). Notably, we found that longer duration of CPR, as well as the dose of epinephrine used during resuscitation, were associated with more severe AGI manifested as more severe hemorrhage and hyperemia (Figure S6F). At 24 hours after ROSC in the vehicle-treated group, obvious necrosis and ileus in small intestine, with gaseous distention in stomach and upper intestine, were found (Figure 8E). Administration of bumetanide partly prevented intestinal necrosis and ileus (Figure 8E).

We performed histological examination with HE staining to further study pathological changes after CA/CPR at 24 hours (Figure 8F). In stomach, there were no obvious changes in forestomach, oppositely, significant damages were found in the glandular stomach: increased erythrocyte diapedesis and interstitial edema in mucosa; interstitial edema, inflammatory cell infiltration, and hemangiectasis in submucosa; inflammatory cell infiltration and thinning in muscularis; occasional ulcerations and necrosis (Figure 8F). In small intestine, including duodenum and the middle section, severe erosion and necrosis were observed: damaged intestinal villi, necrotic ulcerations, increased erythrocyte diapedesis, interstitial edema, inflammatory cell infiltration, hemangiectasis, and thinning in muscularis (Figure 8F). There were only minor histological changes in colon after CA/CPR, including a small number of inflammatory cell infiltration, atrophy of submucosa, decreased number of goblet cell, distortion of the crypt surface, ulceration, and enlargement of glandular cavity (Figure 8F). Treatment with bumetanide alleviated these histological damages caused by CA/CPR (Figure 8F).

To examine the hemodynamic alternations of the most vulnerable intestinal segment, we used hand-held SDF microscope to directly observe the prefusion of mesentery and intestinal wall. In the vehicle-treated CA/CPR mice, there were no signs of obstruction in mesenteric vessels, but significant hypoperfusion in microcirculation of intestinal wall was found, which indicated a manifestation of non-obstructive mesenteric ischemia. Treatment with bumetanide increased the perfusion of intestinal microcirculation compared with vehicle treatment after CA/CPR (Figure 8E).

### Treatment with bumetanide after CA/CPR prevented multiple organ dysfunction in kidney, lung, liver, adrenal gland, and spleen

CA/CPR causes completely whole-body ischemia and reperfusion, it is reasonable to speculate that major organs are at risks to get dysfunction after ROSC. To evaluate the existence/level of post-resuscitation multiple organ injury and effects of bumetanide treatment, we performed histopathological examination of lung, kidney, liver, spleen, and adrenal gland (Figure 9A through 9E).

**Figure 9.**
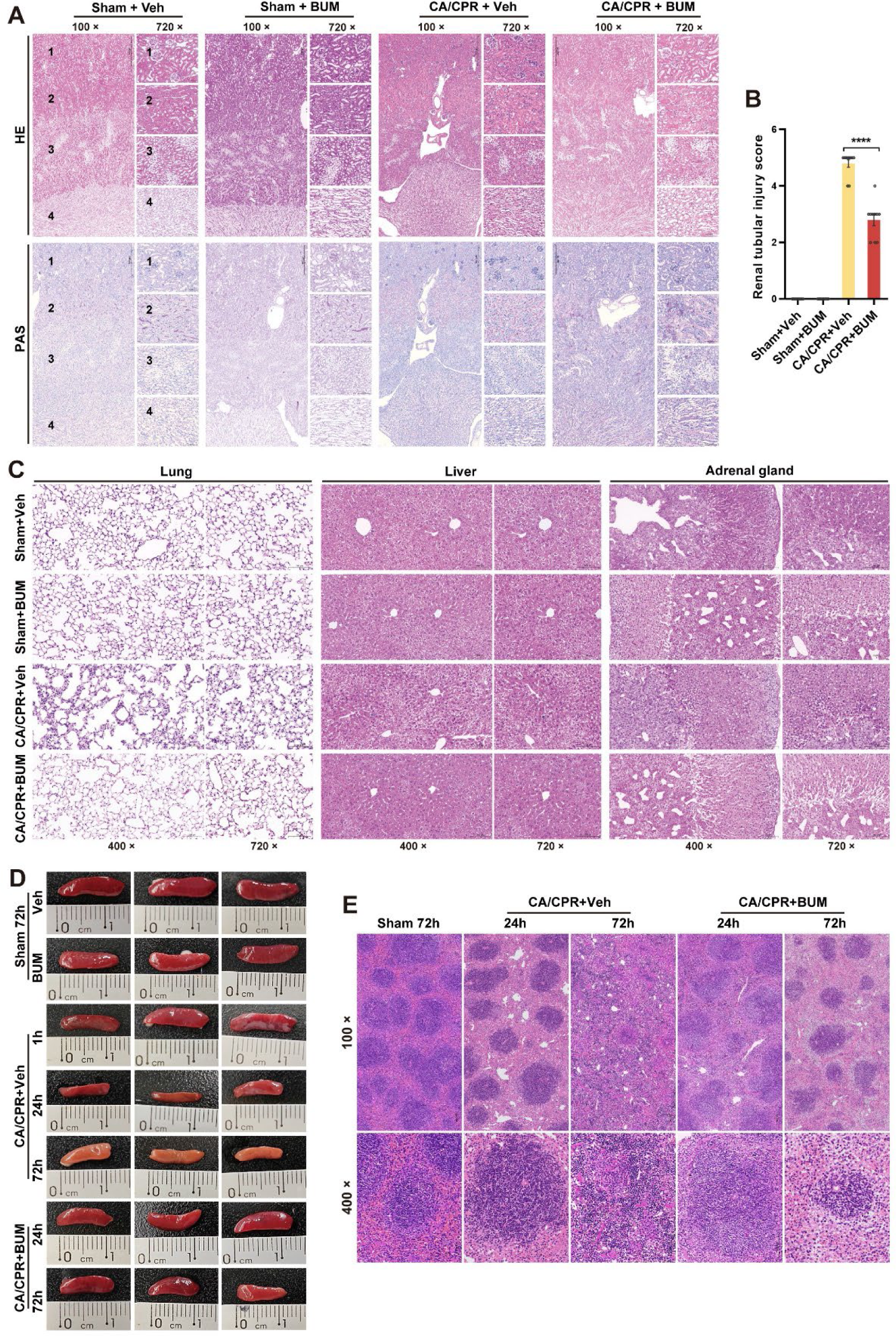
Bumetanide prevented multiple organ injury in kidney, lung, liver, spleen, adrenal gland, and spleen. **A**, Representative photomicrographs of HE-staining and PAS-staining kidney sections 24 hours after CA/CPR. Cross sections and four anatomic regions (1, outer cortex; 2, inner cortex; 3, medulla; 4, pelvis) are shown. CA/CPR caused renal tubular necrosis, inflammatory cells infiltration, interstitial edema, loss of brush border, and increased formation of casts (red blood cell casts, white blood cell casts, pigment casts). Treatment with bumetanide alleviated the histological injuries in kidney. **B**, The renal tubular injury score. (n=10) Data are presented as mean±SEM. \*\*\*\**P* < 0.0001. **C**, Representative photomicrographs of HE-staining lung, liver, and adrenal gland sections at 24 hours after CA/CPR. In lung, CA/CPR caused destroyed alveolars, interstitial edema, thickened alveoli septum, and neutrophil and eosinophil infiltration. In liver, CA/CPR caused inflammatory cell infiltration, cellular edema, spotty necrosis, and fatty degeneration and acidophilic degeneration of hepatocytes. In adrenal gland, obvious inflammatory cell infiltration was found after CA/CPR. Bumetanide treatment reduced above morphological injury after CA/CPR. **D**, Representative photographs of spleen. Atrophy of spleen was not occurred at 1 hour after CA/CPR but was obvious at 24 hours, and bumetanide alleviated CA/CPR-caused atrophy of spleen at 24 hours and 72 hours. The color of spleen faded at 72 hours in the vehicle treated CA/CPR mice. **E**, Representative photomicrographs of HE-staining spleen sections at 24 hours and 72 hours after CA/CPR. Bumetanide alleviated the white pulps injury after CA/CPR.

In kidney, HE staining and PAS staining showed that CA/CPR caused increased urinary casts (red blood cells, white blood cells, pigment casts), acute renal tubular necrosis, loss of brush border, remarkable dilation of renal tubules, infiltration of inflammatory cells, and interstitial edema (Figure 9A). Treatment with bumetanide alleviated the tissue injury as stated above (Figure 9A and 9B).

In lung, CA/CPR induced acute lung injury with thickened alveoli septum, destroyed alveolars, interstitial edema, and increased cell number with inflammatory cell (neutrophil and eosinophil) infiltration, but no bleeding or exudation. Administration of bumetanide mitigated pulmonary histopathological changes (Figure 9C).

In liver, CA/CPR caused markable inflammatory cell infiltration, fatty degeneration of hepatocytes, cellular edema, and disordered hepatic lobule. Additionally, spotty necrosis and acidophilic degeneration could be found in the CA/CPR group. These histopathological findings indicated an acute mild hepatitis occurred after ROSC. The bumetanide-treated mice showed less histopathological changes compared with vehicle-treated mice after CA/CPR, including less spotty necrosis and acidophilic degeneration, which indicated an alleviative hepatitis (Figure 9C).

In adrenal gland, morphological changes after CA/CPR were not obvious except for inflammatory cells infiltration and dilation of vessels in medulla. Decreased number of infiltrated inflammatory cells were observed after bumetanide treatment (Figure 9C).

In spleen, CA/CPR induced an obvious reduction in size (Figure 9D) and obvious pathological injury, including marked damage in white pulps with decreased number of lymphoid follicle and cell numbers, as well as blurred boundary of pulps (Figure 9E). Treatment with bumetanide reduced the loss of size and reduced histological lesions.

Taken together, CA/CPR caused MODS with obvious histological damage in major organ systems, including tissue edema and systemic inflammation with inflammatory cell infiltration in each organ. Treatment with bumetanide prevented MODS and alleviated histological injury with decreased inflammatory cell infiltration and reduced tissue edema.

## Discussion

Here we report that establishing a post-resuscitation MODS model using KCl-induced asystole arrest and successful CPR method in male mice, with manifestations of severe HIBI and EMOD, including dysfunction in heart, lung, liver, spleen, gastrointestinal tract, kidney, adrenal gland, urinary bladder, penis, and spinal cord. Inflammatory cell infiltration and tissue edema could be found in each major organ by histopathological examination after CA/CPR. Systemic administration of bumetanide after resuscitation was safe and dramatically improved outcomes and prevented MODS. The CNS target of bumetanide, NKCC1, has an increased expression immediately after resuscitation, which may represent a novel mechanism of the pathogenesis of PCABI. Neuronal populations in cerebral cortex (especially piriform cortex), amygdaloid nucleus, hypothalamus, corpus striatum, but not hippocampus or thalamus, are selectively vulnerable as confirmed by FJB and HE staining. Pathophysiological mechanisms, including neuronal degeneration, seizures/myoclonus, cerebral inflammation, decreased CBF, BBB disruption, oxidative stress, and brain tissue edema contributed to the progression of PCABI, and treatment with bumetanide alleviated these pathophysiological manifestations. DIA-proteomics analysis of cerebral cortex identified the protein alternations at 24h after CA/CPR. The common DEPs in CA/control may representative the pathogenesis of PCABI, and the common DEPs in the comparison between three groups respectively may have potential to be novel biomarkers of PCABI. We identified NGAL as a novel biomarker for early neuroprognostication at 24 hours after CA/CPR.

In this study, mice had manifestations of severe HIBI after CA/CPR. Seizures may exacerbate brain injury, and myoclonus is the most common type of clinical seizure, which may have a cortical or subcortical origin^2^ that were also vulnerable brain regions in our study. Reduced CBF after ROSC is injurious and directly impairs cerebral oxygen and substrate delivery.^29^ An intact BBB is essential for building and maintaining a microenvironment that allows brain to function properly, and BBB breakdown leads to increased extravasation of immune cells and poorly regulated flux of ions and molecules across the barrier when tight junctions are disrupted and/or transport processes are impaired, generally culminating in neuronal dysfunction, inflammation, and neurodegeneration.^30^ The expression of CNS target of bumetanide, NKCC1, increased immediately after resuscitation. Thus, the alleviation of the foregoing pathophysiological harms by bumetanide treatment was not surprising, given that the preventive benefits of bumetanide by targeting elevated cerebral NKCC1 expression had been well researched.^13,14,17^ We also proved severe cerebral tissue edema after CA/CPR, which is an indication for bumetanide treatment and was reduced as evidenced by histological examinations. The path length for oxygen diffusion from blood into brain tissue may increase due to edema,^29^ and dehydration with bumetanide could optimize brain oxygenation by optimizing oxygen diffusion. The neuroinflammation was severe after CA/CPR and was reduced by bumetanide, which was also observed in other brain injury studies.^18^ Notably, the strong positive brain areas with immunostaining for astrocytes/microglia (like hippocampus) were largely nonoverlapping with FJB-positive areas, which indicated other mechanisms rather than glia activation played a dominant role that mediated neurodegeneration upon PCABI. Previous studies using the same model demonstrated immune cells, including neutrophils and monocytes, but not lymphocytes, infiltration in brain after CA/CPR.^25^ Additionally, we identified increased expression of CD14, the endotoxin receptor, in brain after CA/CPR. Taken together, these results highlighted an infectious complication developed on the basis of sterile inflammation primarily caused by cerebral ischemia-reperfusion injury.

The four key components of the PCAS pathophysiology were identified as PCABI, PCAMD, SIRR, and PPP.^1^ Firstly, PCAMD-induced cardiovascular failure accounts for the most of early death after resuscitation.^2^ Manifestations of PCAMD include hypotension and ventricular systolic/diastolic dysfunction, resulting in reduced cardiac output, arrhythmias, and pulmonary edema.^3,31^ In this study, abnormal but inconsistent electrocardiogram manifestations, as well as histological injury with myocytolysis and myocardial fibrosis, could be found at 24 hours after ROSC. These findings indicated the presence of acute myocardial dysfunction, which reduced the cardiac output and consequent decreased systemic perfusion. The lower organ perfusion limited the oxygen delivery and aggravated the imbalance between oxygen delivery and utilization, leading to secondary ischemia and subsequent organ dysfunction. Administration of bumetanide alleviated the myocardial injury and improved the electrocardiogram manifestations, which indicated an improved heart pump function and optimized perfusion of end organs. Additionally, improved hemodynamics of macrocirculation may also potentially improve microcirculation perfusion. Thus, it is an important mechanism mediated the protective effects of bumetanide on improving outcomes and preventing MODS. Secondly, SIRR was identified as a sepsis-like syndrome, with systemic inflammation, endotoxemia, immunosuppression, and increased risk of infection.^1,31^ In this study, systemic inflammation with inflammatory cell infiltration were found in each major organ system after CA/CPR. Infiltration of eosinophils and neutrophils could be found by histopathological examinations. We proved MODS, including AGI with severe intestinal necrosis, after CA/CPR, and the cerebral proteomic data showed response to external stimulus after CA/CPR, highlighting the presence of infection and consequently a true sepsis. Previous studies using the same model identified the mouse was in an immunosuppressed state after resuscitation, with activation of hypothalamic-pituitary-adrenal axis and atrophy of major immune organs,^25^ which were also found in our study that spleen dramatically atrophied with decreased cell numbers. The bumetanide-treated CA/CPR mice showed less inflammatory cell infiltration in organs and better spleen morphological characteristics, indicating decreased systemic inflammation and preserved immune function for response to external stimulus. Lastly, hyperkalemia represents the PPP that caused CA, and bumetanide natively has kaluretic effects. Thus, bumetanide treatment is beneficial to solve the PPP underlying CA/CPR. Taken together, our study showed that bumetanide could help to solve all four key components of PCAS pathophysiology.

Post-CA MODS is common and associated with mortality, and it is reasonable to postulate that SIRR could affect all organ systems.^8,9^ The progression of post-resuscitation MODS may be driven by hemodynamic dysfunction and oxygenation impairment after resuscitation.^8^ It is difficult to synthesis the pathophysiological framework of post-CA MODS given that multiple molecular and pathological mechanisms and organ crosstalk, through which dysfunction of one organ could lead to the dysfunction of another remote organ, which may make it a vicious cycle and leads to multiple organ failure, even death. This preclinical study provided the evidence that all major organ systems got dysfunction form CA/CPR, which should be verified in future clinical studies. From this view, it is reasonable to close monitor the function and potential progression or undiscovered existence of injury in vital organs. Additionally, optimize organ perfusion is the fundamental treatment strategy,^3,9^ and early prediction, early prevention, early identification, and early intervention for dysfunction/failure in each organ need to be emphasized in clinical management bundles.

The gastrointestinal injury after CA is poorly understood and less studied, but the actual incidence may be surprisingly high.^32^ Ischemic injury, including necrosis, ulceration, mucosal edema or erythema, and non-occlusive mesenteric ischemia, characterized by hypoperfusion without mechanical obstruction, could be found in clinical observations and were significantly associated with mortality.^32,33^ In this study, AGI with severe lesions was obvious after CA/CPR, which appeared as early as soon after resuscitation. The most vulnerable segment was in the middle of small intestine, at the initial part of ileum, characterized by intestinal hemorrhage after operation and necrosis with intestinal obstruction at 24 hours after resuscitation. The presence of black tarry stool also indicated the presence of gastrointestinal bleeding. We identified this most vulnerable intestinal segment showed features of non-occlusive mesenteric ischemia, as evidenced by the medical history, morphological examinations, and focal hemodynamic monitoring. Additionally, we also proved that all gastrointestinal tract was injured after CA/CPR, which highlights the significance of prevention, diagnosis, and treatment of AGI in future clinical practice. The gut was regarded as the motor of critical illness, which was hypothesized to play a central role in the progression of sepsis and MODS mediated by altered formation of toxic gut-lymph, alternations in gut microbiome, and injury of intestinal barrier and consequent bacterial translocation.^34^ And in PCAS, the severity of intestinal injury was associated with endotoxemia,^35^ which indicates the presence of gut-derived infection/bacterial translocation after CA/CPR. Our CA/CPR model showed severe intestinal lesion with transmural necrosis, thus, it is reasonable to speculate the presence of bacterial translocation and endotoxemia, which might lead to sepsis, aggravated systemic inflammation on the basis of ischemia/reperfusion-induced sterile inflammation, and promoted the progression of MODS. Therefore, administration of bumetanide alleviated AGI after CA/CPR, which could decrease the blood endotoxin level and relevant systemic inflammatory responses in theory. However, the exact mechanisms of bumetanide on post-resuscitation AGI protection are still unknown, and whether bumetanide had direct protection effects need to be further investigated.

Acute lung injury/acute respiratory distress syndrome is common and associated with worse outcomes after CA.^36^ Our study showed acute pathological changes in lung and decreased breathing frequency with irregular breathing pattern after CA/CPR. Histological injury with abnormal respiration worsened systemic oxygenation impairment and contributed to secondary hypoxia injury and consequent poor prognosis. Bumetanide alleviated the lung injury and improved breathing after resuscitation, which enables increased delivery of oxygen from pulmonary vasculature to vital organs to alleviate post-resuscitation shock.

Severe acute kidney injury (AKI) occurs frequently after CA, which usually occurs at an early stage after resuscitation and is significantly associated with mortality and poor neurological outcome, and recovery form AKI may be associated with improved survival and good neurological outcome.^37,38^ In the present study, we demonstrated severe morphological renal injury, characterized by tubular necrosis and casts formation, occurred at 24 hours after resuscitation. This severe injury-induced acute renal insufficiency could result in disturbance of water and electrolyte, worsen the homeostasis and remote organ dysfunction. Systemic administration of bumetanide after resuscitation significantly reduced renal morphological injury and relieved system of urinary retention, preventing urinary dysfunction ultimately. Bumetanide is known to have a renal protection activity by increasing renal blood flow, which was mediated by stimulating prostaglandin synthesis and dilating renal vessels.^39^ In this present study, we observed obvious vasoconstriction after CA/CPR, and bumetanide treatment dilated the constricted kidney vessels and alleviated subsequent histological injury.

Post-resuscitation acute liver injury, usually manifested as hypoxic hepatitis characterized by centrilobular cell necrosis and periphery fatty degeneration, is a frequent finding after CA and is associated with mortality.^40^ In our study, acute hepatitis was diagnosed according to pathohistological examination, showing inflammatory cell infiltration, fatty degeneration, and cellular edema of hepatocytes. Spotty necrosis and acidophilic degeneration were found, which are hallmarks of acute mild, but not severe, hepatitis. Taken together, these findings indicate that liver get mildly injured at early phases after resuscitation and may not play an important role in the progression of PCAS in this study.

To the best of our knowledge, this is the first study that performed cerebral cortex DIA-proteomic analysis after CA/CPR. Alternations of cerebral cortex proteins have been well recognized as indicators of pathophysiological changes underlying PCABI, and proteomic investigations have potential to identify pathogenesis and novel biomarkers. Our proteomic data allowed us to define altered proteins upon PCABI, as well as DEPs between poor and relatively good neurological outcomes for early prognostication at 24 hours after resuscitation. The most prominent proteomic changes are related to response to stimulus, metabolic process, regulation of biological process, and endomembrane system in the comparison between CA and control. Immune response and inflammation are prominent in the comparison between poor and relatively good outcomes, which highlights the significance in the progression of PCABI.

We identified 10 potential markers associated with neuroprognostication, and further, we identified NGAL and angiotensinogen are the common DEPs at 24 hours after resuscitation combined with serum proteome profiles from CA patients.^28^ Angiotensinogen, mainly expressed in astrocytes and neurons in brain, is the precursor of angiotensin which is the part of renin-angiotensin system.^41^ Increased cerebral angiotensin may have various effects. For example, angiotensin II showed neuroprotective effects for ischemic stroke through activation of type 2 receptor mediated by vasodilation, and also had neurodegenerative effects by activation of type 1 receptor to promote oxidative stress and inflammatory responses.^41^ LCN2/NGAL is an inflammatory marker produced by a variety cell types, including immune cells (neutrophils, macrophages, dendritic cells), CNS cells (neurons, astrocytes, microglia), and other cells (renal cells, hepatic cells, cardiomyocytes).^42^ The cerebral expression of NGAL under physiological conditions is very low and undetectable, and the function of neuroinflammatory regulation of NGAL is complex with both pro-inflammatory and anti-inflammatory effects.^42^ Systemic inflammation induced by administration of lipopolysaccharide could upregulate NGAL expression in brain.^42^ In the present study, we detected the expression of NGAL by immunofluorescence, and the results showed that expression level is extremely low in the sham groups. Increased expression of expression in piriform cortex, amygdaloid nucleus, hypothalamus, and retrospleninal cortex, but not hippocampus and thalamus, were found after CA/CPR, while post-resuscitation administration of bumetanide decreased the expression in the above areas. We compared the positive expression areas of NGAL and FJB, and found that there were highly overlapped double-positive areas, which indicated the strong and graded correlation between NGAL and neuronal degeneration underlying PCABI. We also detected the increased expression of CD14, which is known as the receptor of endotoxin lipopolysaccharide and acts as a critical early mediator of the innate immune response,^43^ by immunofluorescence. Concordant upregulation of CD14 expression has been revealed in some clinical and pre-clinical disease studies, which suggests the downstream signaling pathway of CD14 might play a role in many pathologies.^43^

There are still some critical knowledge gaps related to the translation of proteomic studies to clinical application. Firstly, it is scarcely possible to obtain brain samples for protein detecting in clinical practice, and serum is the most widely used and most accessible sample for detection of prognostication. Previous clinical studies have shown the serous detectability of NGAL and angiotensinogen,^28^ and a further challenge is focusing on the cerebral-specific quantification of release of these biomarkers. Utilizing invasive neuromoniting to conduct paired sampling of arterial and jugular venous bulb blood makes the quantified transcerebral release of these biomarkers upon PCABI possible,^44^ which allows us to distinguish the cerebral-specific pathogenesis from systemic circulation. Secondly, another important knowledge gap is the accuracy, combination, and optimal threshold levels of these biomarkers for prognostication. Predicting good and poor neurological outcome, with high sensitivity and high specificity, are important for making clinical decisions, and the roles of these identified novel biomarkers in prediction of outcomes require further investigated, elucidated, and experimentally validated. Notably, previous described markers for neuroprognostication after resuscitation, such as neuron-specific enolase, neurofilament light chain, tau, GFAP, and ubiquitin carboxy-terminal hydrolase-1,^45^ were not detected or significantly different in the current study, which need further studies to reveal the reason. Lastly, only cerebral cortex was used for DIA proteomics studies. Our study and previous studies showed neuron subpopulations in different regions are selectively vulnerable, therefore, proteomic alternations in other brain regions or different subregions within cortex are same or heterogeneous need to be validated. Additionally, the nature of DIA proteomic method determines that it only detects quantitative proteomic changes, which lacks information of protein changes in space or at single-cell resolution, as well as information of protein modification (like phosphorylation) and regulation before translating. Novel and advanced methods like spatial omics or multi-omics could provide more detailed information in future studies. CA remains a leading cause of death worldwide, and significant process has been made on improving the rate of ROSC. However, no interventions, during resuscitation or in post-resuscitation care, have been proved to improve long-term outcome, especially survival with intact neurological function.^2,3,5,7^ PCABI is the leading cause of death and disability,^1–3,5,7^ therefore, it is of high priority to understand the pathogenesis and identify biomarkers upon PCABI. It is becoming clear that no single treatment can solve PCABI given that the complexity of pathophysiology, and a multi-pronged approach like bundle-based management strategies is likely required.^6,29^ Optimizing cerebral oxygenation and developing pharmacological neuroprotective strategies remain fundamental and promising approaches to reducing PCABI. Additionally, accurate and early neuroprognostication with novel biomarkers is of great significance for physicians and family to make decisions to avoid futile efforts. Therapies that focus on neuroprotection may compromise extracerebral injured organ systems, therefore, the systemic effects of neuroprotective agents need to be evaluated. From the perspective of translational medicine, we think our findings, that systemic administration of bumetanide is beneficial to PCABI and EMOD that caused by hyperkalemic-CA/CPR, provide significant clinical implications. Post-resuscitation treatment with bumetanide is safe and effective, and these results may expand to other PCAS situations with different CA causes. Using LCN2/NGAL for early prediction of unfavorable neurological outcome at 24 hours after CA/CPR may be a novel approach, and the sensitivity and specificity of which need to be tested in future large-scale clinical studies.

This study has several limitations. Firstly, only young adult male mice were used in this study. The severity of post-resuscitation injury, effects of bumetanide, and proteomic changes underlying PCABI in female, children, and the aged population require further study. Secondly, only KCl-induced asystole arrest model was used in this study. Causes of CA and underlying comorbidities after resuscitation contributes to the heterogeneity of PCAS, therefore, mechanisms of PCABI progression with different causes and therapeutic effects of bumetanide need to be further explored. Thirdly, we were not intended to examine damage in all organ systems after CA/CPR, only major organs were focused in this study. Other organ systems, such as great vessels, thyroid, pancreas, and bone marrow, may also get dysfunction from direct hit by CA/CPR or secondary hit form remote organ injury via organ crosstalk. These damages may also play an important role in the progression of PCAS, and thereby survival and neurological prognostication. Therefore, whether damages in these organ systems are existing and the severity of damage need to be investigated. Lastly, the severity of seizures and effects of bumetanide treatment were not quantified with electroencephalogram. In lack of monitoring equipment, we were unable to make quantitative descriptions of the seizures with the golden standard method, and only clinical observations were performed which partly lacks objectivity. Continuous electroencephalogram-monitoring after CA/CPR could provide more detailed and objective descriptions in future PCABI studies.

In conclusion, the current study showed the presence of severe MODS after CA/CPR, and revealed the safety and protective effects of systemic administration of bumetanide after resuscitation on improving outcomes and preventing MODS. Cerebral-specific quantification of LCN2/NGAL, which has a strong and graded association with pathogenesis of PCABI, could be a novel biomarker for early neuroprognostication at 24 hours after CA/CPR.

## Acknowledgments

We thank SpecAlly Life Technology Co., Ltd. for the help in the analysis of proteomic data.

## Non-standard Abbreviations and Acronyms

BBB: blood-brain barrier
BUM: bumetanide
CA: cardiac arrest
CBF: cerebral blood flow
CNS: central nervous system
COG: cluster ortholog groups
CPR: cardiopulmonary resuscitation
CVC: central venous catheterization
DEPs: differentially expressed proteins
DHE: dihydroethidium
DIA: data-independent acquisition
EMOD: extracerebral multiple organ dysfunction
FJB: Fluoro-Jade B
GFAP: glial fibrillary acidic protein
GO: gene ontology
HE: hematoxylin-eosin
HIBI: hypoxic ischemic brain injury
Iba1: ionized calcium-binding adaptor molecule 1
LCN2: lipocalin-2
LSCI: laser speckle contrast imaging
MODS: multiple organ dysfunction syndrome
NGAL: neutrophil gelatinase-associated lipocalin
NKCC1: sodium-potassium-chloride cotransporter 1
PAS: periodic acid-Schiff
PCABI: post-cardiac arrest brain injury
PCAMD: post-cardiac arrest myocardial dysfunction
PCAS: post-cardiac arrest syndrome
PPI: protein-protein interaction
PPP: persistent precipitating pathology
ROS: reactive oxygen species
ROSC: return of spontaneous circulation
SDF: sidestream dark field
SEM: standard error of the mean
SIRR: systemic ischemia/reperfusion response
Veh: vehicle

